# Chromosomal Aneuploidy in Normal, Non-Neuronal Brain Nuclei of Glioblastoma Patients is Not a Cancer Driver

**DOI:** 10.1101/2025.04.22.649812

**Authors:** Olivia Albert, Shixiang Sun, Jhih-Rong Lin, Moonsook Lee, Chang Chan, Alexander Y. Maslov, Lisa Ellerby, Anita Huttner, Zhengdong Zhang, Jan Vijg, Cristina Montagna

**Author notes:** To whom correspondence should be addressed at Rutgers Cancer Institute, 195 Little Albany St., New Brunswick, NJ, 08901, USA.

## Abstract

Aneuploidy is a hallmark of cancers, including high-grade glioma (GBM), one of the most aggressive brain tumors. To assess whether increased aneuploidy already occurs in normal brain tissue of GBM patients, we performed single-nucleus whole-genome sequencing on 225 non-neuronal cortical nuclei from 12 disease-free individuals and 6 GBM patients, in the latter analyzing both tumor and non-tumor distal regions. Somatic aneuploidy was found in approximately 15% of non-neuronal nuclei in the adult human cortex, with recurrent chromosome 16p aneuploidy in up to 4% of nuclei. In contrast, about 51% of GBM tumor nuclei showed frequent aneuploidy of chromosomes 7 and 10, consistent with known GBM profiles. Notably, non-tumor brain regions from GBM patients exhibited aneuploidy frequencies and patterns similar to controls, including recurrent 16p involvement. These findings indicate that somatic aneuploidy in non-neuronal cells is a normal feature of the adult human brain and not linked to increased GBM risk.

## Main

Aneuploidy, the presence of an abnormal number of chromosomes, results from chromosomal missegregation^1^ and is a hallmark of cancer genomes^2^. By promoting genetic diversity^3^ and conferring selective advantages^4^, aneuploidy facilitates tumor evolution, adaptation to changing microenvironments, and therapy resistance^4,5^. This is particularly relevant in high grade glioma (GBM), an aggressive glial progenitor-derived brain tumor with a median survival of 12-18 months^6^. In GBM, 25-30% of tumors exhibit chromosome 7 gain and chromosome 10 loss, associated with EGFR amplification and PTEN loss^7^. Aneuploidy also occurs in precursor glial lesions, suggesting an early role in gliomagenesis^8^, as observed in premalignant lesions of the colon^9^, gastrointestinal tract^10,11^ cervix^12^, ovary^13^, and breast^14^, reinforcing its possible role in tumor initiation. Somatic mutations in cancer driver genes have been found in multiple normal tissues, in the absence of cancer, such as skin, colon and lung^15^, suggesting that also aneuploidy could be a common feature of phenotypically normal tissues, possibly as an early step in cancer^16^.

Understanding how aneuploidy arises in glial cells is essential to uncover its potential role in GBM initiation. Some studies suggest low-level aneuploidy as an adaptive response to stress^17-19^, while others implicate it in age-related neurodegenerative diseases^20-22^. Discrepancies in findings may reflect differences in methodologies, brain regions, cohort sizes^23-26^, age^27^, and environmental exposures. Here, we studied whether individuals who develop GBM already harbor elevated levels of aneuploidy in glial cells in the brain, predisposing them to tumor-promoting alterations. We performed single-nucleus whole-genome sequencing (snWGS)^28,29^ of phenotypically normal, non-neuronal nuclei from disease-free individuals and GBM patients, with matching non-tumor and tumor tissue for comparison. In tumors, we confirmed frequent gain of chromosome 7 and losses of chromosome 10^7^. In contrast, non-neuronal nuclei from both healthy controls and non-tumor GBM brain regions showed low aneuploidy frequencies, with recurrent involvement of chromosome 16p. These findings suggest that aneuploidy is a normal feature of non-neuronal brain cells, but unlike other mutations, is not a driver of gliomagenesis.

## Results and Discussion

To assess aneuploidy in non-neuronal nuclei of the adult, normal human brain, we first analyzed disease-free cortical tissue from 12 healthy individuals (aged 51.8±13.8 years **Extended Data Table 1**) using low-coverage-single nucleus whole-genome sequencing (snWGS). Nuclei were isolated from frozen brain samples using Fluorescence-Activated Cell Sorting (FACS) after NeuN and DRAQ5 staining (**Supplementary Figure 1A**). This strategy enabled detection of euploid (2n), polyploid (3n), and aneuploid nuclei, including chromosome arm and whole-chromosome alterations (**Supplementary Figure 1B**). This strategy was validated using euploid and Trisomy 21 (T21) control cell lines (**Supplementary Figure 1C-G, Extended Data Tables 1-4**).

Only autosomes were analyzed; sex chromosomes were analyzed separately due to their distinct biology^30^. The mean sequencing depth for healthy control nuclei (HC) was 0.15 ± 0.13X (**Extended Data Table 5, Supplementary Figure 2A**). Among 105 NeuN-neg HC nuclei, 89 (84.8%) were euploid and 16 (15.2%) were aneuploid (**Figure 1A**). Of the aneuploid nuclei, 10 (9.5%) had chromosome arm aneuploidies, 1 (0.9%) showed an arm level change in a triploid background, and 3 (2.9%) had both arm and chromosome-level alterations in a diploid background. Two nuclei (1.9%) were triploid without detectable aneuploidies (**Figure 1B-1C**). Gains and losses occurred at similar frequencies, accounting for 10 (35.7%) and 18 (64.3%) aneuploidies, respectively (p=0.094, chi-square test) (**Supplementary Figure 2B**). Each aneuploid nucleus harbored either gains or losses, never both (**Supplementary Figure 2C**), and no chromosome-specific bias was observed (p=0.81) (**Supplementary Figure 2D**).

**Figure 1:**
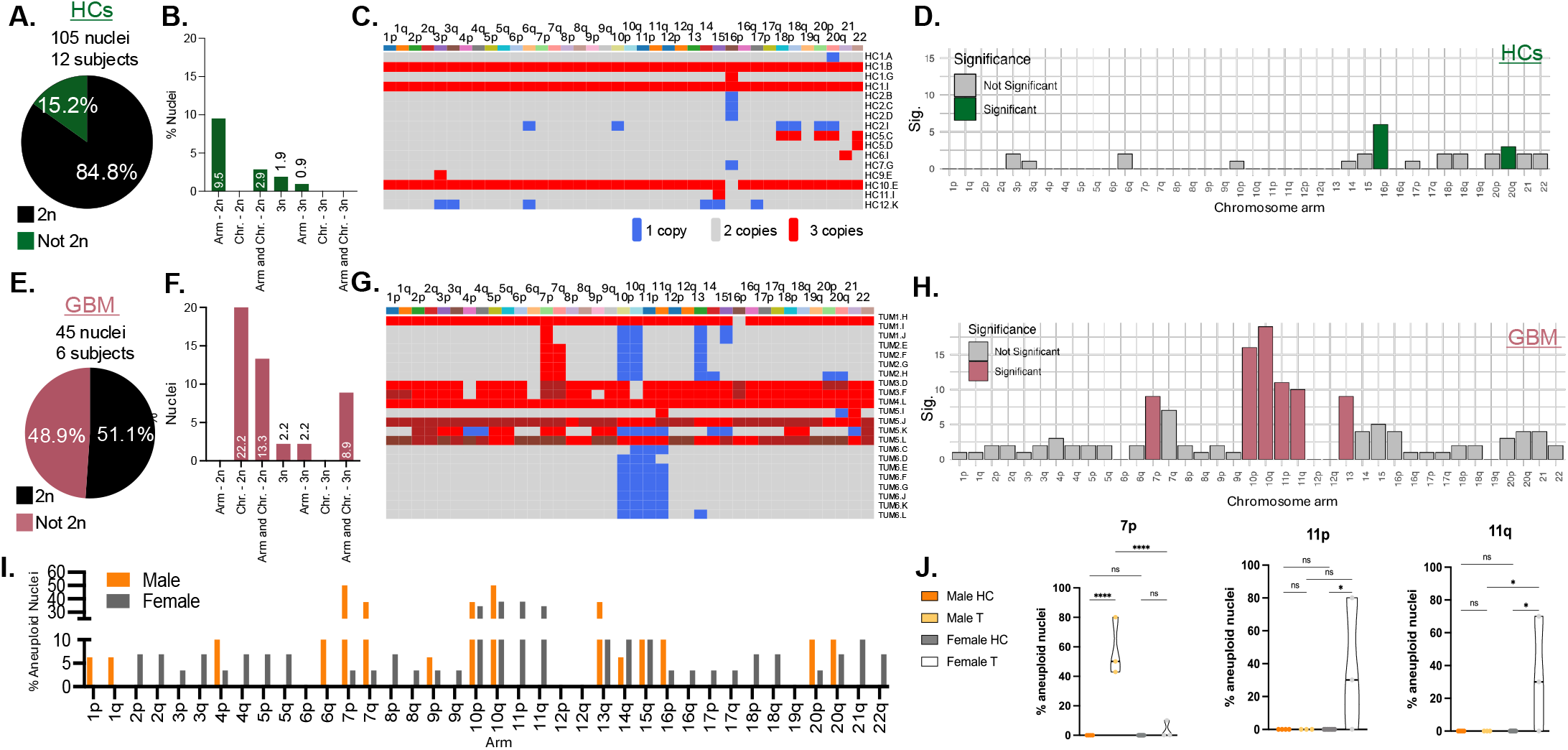
Chromosomal aneuploidy in NeuN-negative cells of healthy and GBM cortex. **A**. Proportion of euploid (black) and aneuploid (green) nuclei from healthy control brains. **B**. Distribution of aneuploidy types in healthy controls: diploid (2n) nuclei with chromosome-arm level aneuploidy, 2n with whole-chromosome aneuploidy, 2n with both arm- and chromosome-level aneuploidy, triploid (3n) without aneuploidy, and 3n with chromosome-arm level aneuploidy. Numbers inside bars indicate the respective percentages. **C**. Heatmap showing chromosome copy number states in aneuploid nuclei from healthy controls (euploid nuclei excluded). X-axis: chromosomes; Y-axis: individual aneuploid nuclei. Blue, grey, and red indicate one, two, or three copies, respectively. **D**. Monte Carlo analysis identifying chromosomes and chromosome arms significantly more aneuploid than expected by chance. Significant chromosomes and arms are shown in green; non-significant in grey. Y-axis: statistical significance; X-axis: chromosomes and chromosome arms. **E**. Proportion of euploid (black) and aneuploid (pink) nuclei in GBM tumor tissue. **F**. Distribution of aneuploidy types in GBM tumor nuclei (TUM), categories as in panel B. **G**. Heatmap of aneuploid tumor nuclei (euploid nuclei excluded). X-axis: chromosomes; Y-axis: individual aneuploid nuclei. Significant chromosomes shown in pink. **I**. Percentage of aneuploid nuclei in GBM tumors, stratified by sex (orange: males; grey: females). **J**. Chromosomes showing significant sex-specific differences in aneuploidy: 7p, 11p, and 11q. Male and female tumors compared to controls. 7p: Male HC vs. Male TUM (p=<0.0001); Male TUM vs. Female TUM (p=<0.0001), 11p: Female HC vs. Female TUM (p=0.0236); Male TUM vs. Female TUM (p=0.0599), 11q: Male TUM vs. Female TUM (p=0.0475), Female HC vs. Female TUM (p=0.0178).

In healthy controls, 24 chromosome arms exhibited no aneuploidy, while 12 (3p, 3q, 6q, 10p, 14, 15, 17p, 18p, 18q, 20p, 21, and 22) showed aneuploidy in ≤2% of nuclei. Chromosomes 16p and 20q displayed significantly higher aneuploidy frequencies compared to random chance (16p: p=1.21×10^−4^, 20q: p=**0.0417)** (**Figure 1D**). Four nuclei, one each from HC1, HC3, HC5 and HC7, had a 2.03Mb segmental loss (chr16:14,817,633-16,845,164) (**Extended Data Figure 1A**). Gene enrichment analysis showed a significant association with 16p13.11 copy number variation syndrome (WP5502; adjp 4.888×10^−19^) (**Extended Data Table 6**), linked to neurodevelopmental disorders, including intellectual disability, autism, schizophrenia, epilepsy, and Attention-Deficit/Hyperactivity Disorder (ADHD)^31^.

The percentage of aneuploid nuclei per individual ranged from 0 to 44% (**Supplementary Figure 2E**), with no significant difference in overall genome alterations **(Supplementary Figure 2F)**. Notably, euploid and aneuploid groups differed significantly in their euploid cell counts (W = 25.5, p = 0.02433) and in the number of affected chromosomes (W = 27, p = 0.0095), highlighting variability within healthy controls (**Extended Data Figure 1B)**.

To quantify chromosomal instability, we applied three metrics: structural score (number of ploidy changes for each chromosome), aneuploidy score (number of gained or lost chromosomes per subject), and heterogeneity score (variability of aneuploidies among nuclei from the same subject). Aneuploidy scores were similar across most individuals, with the exception of HC1 and HC10, which had higher scores due to the presence of 3n nuclei (two in HC1, one in HC10). Individuals with scores near zero were excluded from the plot (**Extended Data Figure 1C**).

We then analyzed both tumor (TUM) and non-tumor (NT) brain regions from six GBM subjects (mean age 63 ± 5.8) (**Extended Data Table 7**). The average sequencing depth was 0.21 ± 0.34 for TUM and 0.13 ± for NT (**Extended Data Table 8** and **Supplementary Figure 3A**).

Of 45 TUM nuclei, 23 (51.1%) were euploid, and 22 (48.9%) were aneuploid (**Figure 1E**). Among the aneuploid nuclei, 10 (22.2%) exhibited chromosome-level aneuploidies, 6 (13.3%) showed arm- and chromosome-level aneuploidies, 1 (2.2%) was triploid without aneuploidies, 1 (2.2%) had arm-level aneuploidy in a triploid background, and 4 (8.9%) harbored arm- and chromosome-level aneuploidies in a polyploid background (two in a 3n background and two in either a 4n or 5n background) (**Figure 1F-G** and **Extended Data Table 9**). Chromosome arms with aneuploidy in ≤11% of nuclei included 1p, 1q, 2p 2q, 3p, 3q, 4p, 4q, 5p, 5q, 6q, 8p, 8q, 9p, 9q, 14, 15, 16p, 16q, 17p, 17q, 18p, 18q, 20p, 20q, 21 and 22. Chromosome arms 7p, 10p, 10q, 11p, 11q, and 13 were significantly more aneuploid than expected by chance (p-values: 9.77 × 10−^3^ for 7p, 2.63 × 10−6 for 10p, <2.63 × 10−6 for 10q, 7.89 × 10−4 for 11p, 2.96 × 10− ^3^ for 11q, and 9.77 × 10− ^3^ for chromosome 13) (**Figure 1H**). Overall, 35 (25%) chromosome arm gains and 105 (75%) losses were identified, with no significant differences between gains and losses (p=0.0922) (**Supplementary Figure 3C-D**).

GBM incidence is 1.6 times higher in men than in women^32,33^. Here, sex-specific differences were also observed (**Figure 1I**), with chromosome 7p significantly more aneuploid in male TUM (p=<0.0001) compared to female and chromosome arms 11p (p=0.0599) and 11q (p=0.0475) either near significant or significantly more aneuploid in female TUM (**Figure 1J**) compared to male. Whole-genome duplication (WGD), common in tumors^34^, was significantly associated with aneuploidy in tumors (p = 0.0013; **Extended Data Table 10**). WGD nuclei were 18 times more likely to be aneuploid than diploid ones (odds ratio 18.08, 95% CI: 2.15– 849.90).

In NT nuclei, 41 (80.4%) were euploid and 10 (19.6%) were aneuploid (**Figure 2A**). Among aneuploid nuclei, 7 (13.5%) had arm-level alterations, 1 (1.9%) had both arm- and chromosome-level changes in a diploid background, 1 (1.9%) had arm-level aneuploidy in a triploid background, and 1 (1.9%) had both in a triploid background (**Figure 2B-C**). Of the 51 NT nuclei, 19 chromosome arms showed no alterations. Eighteen arms (4p, 5p, 8q, 9p, 10p, 11p, 11q, 12p, 12q, 13, 14, 15, 16q, 17q, 20p, 20q, 21, and 22) were altered in < 5% of nuclei. Aneuploidy on 16p was observed in 14% of nuclei and was significantly altered **compared** to random chance (p = 7.89 × 10^−6^, **Figure 2D)**. Among all NT aneuploidies, 5 (17.8%) were gains and 23 (82.4%) were losses. Each aneuploid nucleus had either gains or losses; only one had both (**Supplementary Figure 3E-G**).

**Figure 2:**
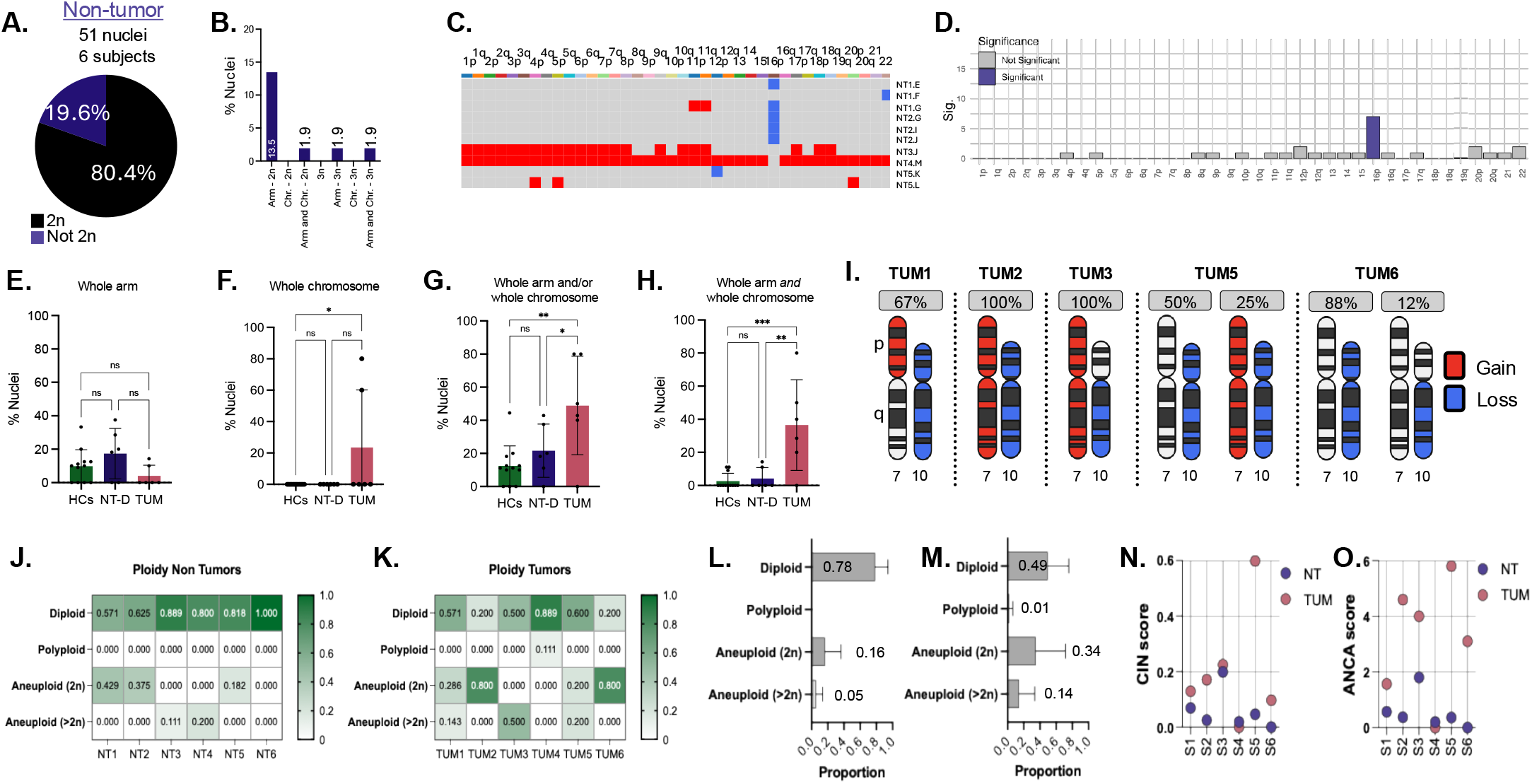
GBM aneuploidy signature detected in tumors, not matching non-tumor regions. **A** Proportion of euploid (black) and aneuploid (purple) nuclei in the non-tumor-(NT) cohort. **B**. Distribution of aneuploidy types in NT: diploid (2n) with chromosome arm-level aneuploidy, 2n with whole-chromosome aneuploidy, 2n with both arm- and chromosome-level aneuploidy, triploid (3n) without aneuploidy, and 3n with arm-level aneuploidy. Numbers inside bars indicate percentages. **C**. Heatmap showing chromosome copy number states in aneuploid nuclei from non-tumor regions (euploid nuclei excluded). X-axis: chromosomes; Y-axis: individual aneuploid nuclei. Blue, grey, and red indicate one, two, or three copies, respectively. **D**. Monte Carlo analysis identifying chromosomes and chromosome arms with significantly more aneuploid than expected by chance in non-tumor tissue. Significant chromosomes and arms are shown in purple; non-significant in grey. Y-axis: significance; X-axis: chromosome arms. **E-H**. Bar graphs comparing the percentage of aneuploid nuclei across healthy controls (HC), NT, and tumor (TUM) cohorts. Y-axis: percentage of nuclei; X-axis: cohort. **E**. Arm-level aneuploidy; **F**. Whole-chromosome aneuploidy (HCs vs. TUM p=0.0419); **G:** Combined arm-level and or whole-chromosome aneuploidy (HCs vs. TUM p=0.0023, NT vs. TUM p=0.0499); **H**. Composite category including all aneuploidy events (HCs vs. TUM p=0.0003, NT vs. TUM p=0.0021). Statistical comparisons performed by One-way ANOVA. **I**. Chromosome arm-level summary of tumor-specific aneuploidy, focused on chromosomes 7p/q and 10 p/q. Each GBM tumor is represented by one or two sets of chromosome arms, depending on the presence of co-occurring aneuploidy patterns. Gains are shown in red, losses in blue, and euploid in black and white,. **J-K**. Ploidy state distributions in NT (**J**) and TUM (**K**) regions by subject. X-axis: GBM subject; Y-axis: ploidy state. The number inside the squares indicate the percentage. **L-M**. Summary plots of ploidy states in NT (**J**) and TUM (**K**) nuclei; X-axis proportion; Y-axis: ploidy state. Number inside bars indicate the proportion plotted for each. **N**. Chromosomal instability (CIN) scores for NT (purple) and TUM (pink) regions. X-axis: GBM subjects; Y-axis: CIN score. **O**. Average copy number alterations (ANCA) for NT (purple) and TUM (pink) subjects. X-axis: individual GBM subjects; Y-axis: ANCA score. A significant difference in ANCA score was observed (t(10) = −2.97, p = 0.014).

Chromosome instability metrics showed distinct patterns for individual subjects, indicating higher instability in some GBM cases (**Extended Data Figure 2A**). NT regions clustered in the lower left of the plot, indicating low chromosomal instability (**Extended Data Figure 2B**). Euclidean distances between NT and TUM differed significantly (p = 0.03), highlighting the differences between these regions (**Supplementary Figure 3H**).

No significant differences in whole-arm or whole-chromosome aneuploidy were found between HCs and NT (**Figure 2E-F**). Tumors had significantly more whole-chromosome aneuploidy than HCs (p=0.0023; **Figure 2F**). Nuclei with either arm or chromosome-level changes were more frequent in tumors than in HCs (p=0.0023) and NT (p=0.0499; **Figure 2G**). Nuclei with both were also significantly enriched in tumors (HCs versus TUM: p=0.0003; NT-D v TUM: p=0.0021; **Figure 2H**).

To identify tumor-specific chromosomal alterations, we compared matched tumor and non-tumor nuclei (**Supplementary Figure 3I-J**). Gains of chromosomes 7 and loss of chromosome 10 were present in 30 to 40% of tumors (**Figure 2I**) and absent in every NT nucleus. Although some subjects’ NT and TUM regions had similar aneuploid cell proportions and CIN scores (**Figure 2 J-N)**, all tumors had significantly higher Average Number of Copy Number Alterations (ANCA) scores than their matched non-tumor tissues (t(10) = −2.97, p = 0.014), with the mean ANCA score of 0.57 in NT and 3.23 in TUM (95% CI: −4.67 to −0.66; **Figure 2O**).

Loss of X (LoX) was detected in 10.7% of all female nuclei (n=107), one from a healthy control female and nine in tumors. Tumor 4 had the highest LoX frequency (66%) but the fewest autosomal aneuploidies, suggesting that X loss may impair function independently. LoX was not observed in males. All X chromosome gains involved the Xp arm, and X chromosome gains were randomly distributed across groups, whereas X loss was specific to female tumors (**Supplementary Figure 4**).

## Conclusions

Here we employed single-nucleus whole-genome sequencing to examine whether elevated aneuploidy is a defining feature of non-neuronal cells of normal brain tissue distal from tumors in individuals with high grade glioma, potentially predisposing them to tumor development. While aneuploidy frequencies in glia of the normal human brain have never been reported, there is considerable controversy about aneuploidy in human neurons. Early studies using FISH reported aneuploidy frequencies ranging from 5.2% to 11.8% for chromosome 21^35,36^. However, these findings were later disputed by other authors using single-cell sequencing assays, suggesting that aneuploidy frequencies were very low^37-39^ and show a much lower frequency of oncogenic variants^40^.

To establish a baseline aneuploidy landscape in non-neuronal cells of the adult human brain, which has never been described, we first analyzed NeuN-negative cells from healthy control subjects. Analyzing both whole-chromosome and chromosome arm-level aneuploidies, our analysis identified a total aneuploidy frequency of 15.2% in healthy subjects. Since aneuploidy is based on mitotic errors our results on mitotically active glial cells, also using single-cell sequencing, may not be that surprising, yet our results provide the first systematic foundation for comparative studies of aneuploidy in relation to brain cancer.

An unexpected finding in healthy controls was the recurrent segmental copy number alteration observed in the healthy adult human brain, mapping to chromosome 16p11.2. This locus is associated with neurodevelopmental disorders, as deletions of 16p11.2 increased the risk for autism spectrum disorder (ASD) and cognitive impairment, while duplications are linked to ASD and schizophrenia^31^.

In non-neuronal cells from GBM tumors, aneuploidy frequency considerably higher and included characteristic GBM aneuploidies - gains of chromosome 7 and losses of chromosome 10. Interestingly, our data reveal striking sex-specific patterns of aneuploidy in GBM. Gain of 7p was markedly more frequent in male tumors (53%) than in female tumors (7%), suggesting it may confer a selective advantage in male gliomagenesis. In contrast, female GBM tumors exhibited increased frequencies of 11p and 11q. These result highlight potent sex-biased mechanisms of chromosomal instability in GBM and underscore the need to investigate sex-specific genomic drivers of glioma progression.

Non-neuronal cells from normal brain tissue distal to the tumor in GBM patients showed an aneuploidy landscape similar to that of healthy controls, including the frequent involvement of chromosome 16. Notably, hallmark GBM aneuploidies involving chromosomes 7 and 10 were absent in patient-matched non-tumor tissue, suggesting that aneuploidy in histologically normal brain does not inherently predispose cells to malignant transformation.

## Acknowledgements

Research reported in this publication was supported by AG068908-02 to Cristina Montagna, the late Dr. Judith Campisi, and Dr. Lisa Ellerby. We thank the Molecular Cytogenetics Core at Albert Einstein College of Medicine, particularly Dr. Jidong Shan and Dr. Yinghui Song, for their assistance with this study. Services, results, and/or products supporting this research were provided by the Biospecimen Repository and Histopathology Service Shared Resource at the Rutgers Cancer Institute of New Jersey, supported in part by funding from NCI-CCSG P30CA072720-25. Additional support was provided by the Einstein-Sinai Diabetes Research Center (P60DK020541). All patient samples were obtained from the NIH NeuroBioBank.

## Material and Methods

### Subjects Cohort and Tissue Collection and Processing

Post-mortem human brain tissues were obtained from the NIH NeuroBioBank (NBB)^41^. All experimental procedures were approved by the Internal Review Board of the Albert Einstein College of Medicine (IRB #2018-9792). To establish a cohort of both GBM and healthy subjects, samples from the NBB were filtered to identify GBM patients with tumors preserved as frozen tissue based on experimental requirements. This process identified 6 GBM subjects (age: 63 ± 5.8 years; 3 females and 3 males). For each GBM subject, tissue was obtained from the tumor core and a non-tumor region distal from the tumor. To generate a control cohort, we selected age-matched healthy controls with brain regions matching, or as close as possible to, the non-tumor regions of the GBM cohort (age: 50.4 ± 12.4 years; 7 females and 5 males) (**Extended Data Table 1 and 7)**. Control samples were further filtered to exclude subjects with co-morbidities known to impact brain function (i.e., diabetes, Alzheimer’s). Approximately 100mg of cryopreserved tissue was obtained for each subject. All tissues were reviewed a board-certified neuropathologist, Dr. Huttner, who confirmed the GBM diagnosis in the tumor specimens and verified that non tumor tissues were free of tumor cells based on H&E staining. GBM tumors displayed hallmark histological features, including high cellularity, pleomorphism, microvascular proliferation, and pseudopalisading necrosis.

### Nuclei isolation

The nuclei isolation processes were adapted from Corces et al^42^ and optimized for frozen brain tissues. While the original method used a dounce and pestle homogenizer, this approach left residual tissue fragments, limiting nuclei extraction. To address this, tissue dissociation was performed using the POLYTRON PT 1200 E (Kinematica, 11010025 Lucerne, Switzerland) which improved efficiency. A 2 mm biopsy punch core (Integra 33-31-P/25, Preston NJ) was used to collect tissue fragments from the frozen brain, which were transferred to a 50 mL tube containing 2 mL of cold 1x Homogenization Buffer (sucrose, EDTA, 10% NP-40, PMSF and β-mercaptoethanol) and allowed to thaw on ice for approximately five minutes. Tissues were homogenized using the POLYTRON PT 1200 E set at half speed for 1.5 minutes on ice. An Iodixanol (Sigma D1556-250ML, St Luis MO) gradient was prepared in a 15 mL tube by layering a 29% Iodixanol solution (sucrose, PMSF, β-mercaptoethanol, CaCl2, Mg (Ac)2 and Tris pH7.8) over a 35% iodixanol solution containing the same reagents. The brain homogenate was mixed with a 50% Iodixanol solution (containing the same reagents) in a 1:1 ratio and layered on top the 29% solution. Clear separation of layers was observed at 3 mL and 6 mL within the 15 mL tube, indicating the formation of a gradient. The gradient was centrifuged at 3,000 RCF without brake for 20 min at 4°C in a pre-chilled centrifuge (ThermoFisher 75009509, Waltham MA). After centrifugation, nuclei were isolated form the interface between the 29% and 35% layers and carefully collected for downstream experiments.

### FACS sorting of cortical NeuN-negative single nuclei

Isolated nuclei were incubated in PBS containing 10% goat serum (ThermoFisher 50197Z, Waltham MA) for 1 hour on ice to block non-specific binding. After centrifugation at 4,000 RPM for 10 minutes, the nuclei were resuspended in PBS containing NeuN Alexa-488 antibody (ThermoFisher MAB377X, Waltham MA) at a 1:500 dilution and kept on ice for 1 hour. Following this, nuclei were then spun down again, resuspended in PBS with DRAQ5 DNA dye (eBioscience 65-0880-92, San Diego CA) at a 1:500 dilution, filtered through a sterile CellTrics 20 *μ*m disposable filter (Sysmex 04-004-2325, Lincolnshire IL) into a 5 mL Polystyrene Round-Bottom tube (Corning Falcon 352058, Corning NY), and kept on ice until sorting. NeuN-neg DRAQ5+ single nuclei were sorted Using the MoFlo Astrios Cell Sorter equipped with a 100 *μ*n nozzle with a sheath pressure of 25 psi, into 12 PCR Tube Strips 0.1 mL + Cap strips, flat bottom (Eppendorf 0030124820, Hamburg, Germany), each containing 3 *μ*L PBS. Following isolation into single PCR strips, nuclei were capped, centrifuged, immediately placed onto dry ice, and stored at −80 ^°^C until further use. Prior to FACS, antibody dilutions were optimized using varying concentrations and nuclei were dropped on slides for visualization under a fluorescence microscope to confirm specific immunofluorescence staining.

### Ultra-low coverage single nuclei whole genome sequencing

Single nuclei libraries were generated for single NeuN-neg nuclei using the PicoPLEX Gold Single Cell DNA-seq kit (Takara R300669, San Jose CA), and indexed using DNA single index Kit (Takara R400660, San Jose CA). To minimize PCR-induced errors, amplification cycles were limited to 8. Library quality (size of DNA fragments and regional molarity) was assessed via Tapestation, and those with a wide peak centered around 800bps and a regional molarity greater than 100 pmol/l were deemed suitable for sequencing. Sequencing was performed on NextSeq500 in the Genomics Center at Rutgers New Jersey Medical School using the 2×150 bp sequencing mode. Raw sequence reads were adapter- and quality-trimmed using Trim Galore (version 0.4.1) and aligned to GRCh37-hg19 using BWA MEM (version 0.7.13)^43^. PCR duplications were removed using samtools (version 0.1.19)^44^ Reads were realigned around the known INDELs, and base qualities were recalibrated based on known SNPs with GenomeAnalysisTK (version 3.5) (https://gatk.broadinstitute.org/hc/en-us). The reads with mapping quality above 30 were kept for downstream analysis. Aligned bam files were converted to bed files using bamToBed (version 2.30.0)^45^. Ginkgo was used for identifying copy number variations, using 500K bin size and all other settings as default^46^.

Control samples, including a euploid (Coriell GM12878, Camden NJ) and a Trisomy 21 cell line (Coriell AG08942, Camden NJ) were sequenced in each sequencing round to verify sequencing accuracy and establish a baseline noise inherent to this single-cell sequencing approach.

Sequencing coverage was assessed by calculating the proportion of bases covered for each chromosome relative to chromosome length. Uniformity of coverage was verified before analysis. Regions with reduced coverage due to sequencing challenges, such as telomeres, centromeres, and GC-rich regions, were blacklisted as low-confidence regions and excluded from further analysis.

### Statistical Analyses

One-way ANOVA was used to assess significance on bar charts comparing aneuploidy frequencies across multiple cohort groups, while t-test was used for comparison between two groups. For bubble plots, Euclidean distance measurements were calculated, and one-way ANOVA was performed to compare the distances across the cohort groups.

### Simulation-Based Threshold Determination for Significant Deviations in Chromosome Copy Number Variation

To estimate expected non-diploid events per chromosome in each cohort, we performed 10,000 simulations using the number of analyzed nuclei for each group, assuming each nucleus contained 38 chromosomes and chromosome arms. The set.seed(123) function in R was used for reproducibility of random number generation. Non-diploid counts were simulated using a binomial distribution, with the probability of success set to 1/38 per chromosome. A significance threshold, set at the 90^th^ percentile of the simulated counts, was used to identify chromosomes with significant deviations. Chromosome counts were reshaped using the pivot_longer function from the tidyr package for individual chromosome analysis. The data was grouped by chromosome, and counts were calculated based on whether the chromosome count was equal to 2 (diploid) or not. After summarization, the grouping was removed, and total counts were compared against the established threshold.

### Group comparison

Healthy controls were classified based on their chromosomal status into euploid and aneuploid groups based on the counts of aneuploid cells in 2n and 3n categories, and the number of affected chromosomes. Subjects were designated as euploid if they had no aneuploidy detected in any of the measured categories (Arm-2n = 0, Arm & Chr. 2n = 0, Arm-3n = 0, Arm & Chr. 3n = 0 and 3n = 0); otherwise, they were classified as aneuploid. The Mann-Whitney U test (Wilcoxon rank sum test) was performed to assess differences between the euploid and aneuploid groups for each relevant variable, employing normal approximation for cases with ties. Statistical analyses were conducted using R version [RStudio 2024.09.0], and a significance level of p < 0.05 was considered statistically significant.

### Gains versus losses NT and TUM

We evaluated chromosomal gains, losses, and no copy number change across 96 single cells (51NT and 45TUM) spanning 38 chromosomes and chromosome arms. Each segment was classified as gain, loss, or no change based on deviations from the modal chromosomal copy number (2n). To determine whether gains or losses were more likely to occur, we performed an exact binomial test on the total counts of gains and losses across all nuclei. The null hypothesis assumed an equal probability (50%) for gains and losses. Out of 3,648 total informative events (excluding unchanged segments), 309 were gains and 71 were losses, with 3,268 segments showing no change. The test was performed using the binom.test() function in R, with the alternative hypothesis that the probability of gain differs from 0.5.

## Data Availability

Raw sequencing data (FASTQ files) have been uploaded to The NeuroBioBank Data Repository within the NIMH Data Archive (NDA) under project ID C3917 “Whole-Genome Sequence Analysis of Postmortem Human Brains from the NIH NeuroBioBank”. The data will be made available upon acceptance of the manuscript. Additionally, metadata associated with the dataset, including (e.g., sample information, gender, disease state), can be accessed through the same repository.

